# A water-reward task assay for evaluating mouse mutualistic cooperative behavior

**DOI:** 10.1101/2021.02.06.430037

**Authors:** Weixi Feng, Yanli Zhang, Ze Wang, Tianqi Wang, Yingting Pang, Ying Zou, Huang Huang, Chengyu Sheng, Ming Xiao

**Author notes:** Corresponding to C.S.,; M.X.,. These authors contributed equally: Weixi Feng, Yanli Zhang.

## Abstract

Social cooperation is fundamentally important for group animals but rarely studied with mice because of their natural aggressiveness. In the present work, we induced pairs of mice to develop a mutualistic cooperative behavior in a non-divided chamber. Each mouse was first trained to learn to use a water dispenser by occupying a particular zone served as a switch to the dispenser. Two trained mice were then put into a chamber containing two separate zones jointly controlling two dispensers. We recorded the latency before each co-drinking, the number and cumulated time of co-drinking each day during the test. These parameters served as quantitative measurements of cooperative behavior in mice. The whole procedure includes preparation, training and testing phases, which take 15 days in total. This assay provides detailed procedures and analytical methods for investigators to characterize and quantify the mutualistic cooperative behavior. The use of mice as subjects allows convenient coupling to other behavior assays and is amiable to genetic manipulations for mechanistic study.

## Introduction

Cooperation is defined as two or more individuals working together to achieve a common goal. Cooperative behavior increases participants’ survival fitness and thereby promotes the species’ ecological success^1^. Cooperation can be briefly categorized into two groups: mutualistic cooperation, in which participants gain immediate benefits, and reciprocal altruism, where one or some participants benefit less or even have net costs at the time of behavior but get compensated later^2-4^. Reciprocal altruism is pervasive in human society but is only sparsely reported in nonhuman animals, while mutualistic cooperation is widely evidenced across various animal species^5^. For reciprocal altruism, recent fMRI studies on human subjects identified important brain regions, such as the caudate nucleus and the orbitofrontal cortex in reward processing^6-8^; on the other side, although seeming simpler and more experimentally tractable, the neuronal and molecular mechanisms beneath mutualistic cooperation remain unclear. This calls for the development of animal models and suitable behavioral assays that could reliably assess animal’s cooperative ability.

Cooperation is a complex behavior with an intrinsic social dimension and at the same time requires certain levels of cognitive abilities. Many experimental studies are conducted with nonhuman primates and dolphins, in which animals are expected to pull a rope or handles or push buttons^9-11^. Rats are prosocial animals and are found to show generalized altruistic behavior, making it another excellent model for studying cooperative behavior^2,12^. The mouse model is extensively employed in various social and cognitive behavior studies, but its use in cooperation research is much sparse. In a recent study, Il-Hwan et al. employed male C57BL/6J mice as subjects; with wireless deep-brain stimulation into the medial forebrain bundle as a reward, they found mice could develop and observe a ‘reward zone allocation’ rule and displayed robust positive reciprocity towards their partners^13^. This work demonstrates mice can develop higher-level cooperative resolution of conflicts in some cases. The mouse model is superior in the rich genetic resources and tools, established behavior-tracking techniques, and the vast knowledge gained from mouse neurobiology work. Therefore, developing cooperative behavior assays using the mouse model is particularly attractive for mechanistic study.

In the present work, we established a new water-reward assay to investigate mutualistic cooperative behavior in mice. This paradigm required mice to be first trained in a chamber individually, where it learns to switch on a water dispenser for drinking by occupying a particular zone. Then, two trained mice were paired and put into a chamber containing two water dispensers and two zones. They could get water together only when they occupy both zones simultaneously, a behavior we regard best explained by mutualistic cooperation. Mice were water-deprived 8 hours before each training or testing to enhance their motivation for the task. During the testing period, we observed a consistent reduction in co-drinking latency, and a consistent increase in co-drinking number and cumulated co-drinking time each day, together reflecting a gradually enhanced cooperative performance, as stochastic co-occurrence should predict flattened curves through time.

### Comparison with other methods

Our cooperative water-reward assay is conceptually similar to the synchronized button-pressing task for bottlenose dolphins^10^, or the coordinated platform-locating task for Lister Hooded rats^12^, in which fish or food pellets served as a reward, respectively. However, using house mice as subjects allows easier molecular-genetic manipulations and convenient coupling to other established behavior assays. In a recent study, Kyung et al. used *Shank2* KO and *Shank3* KO mice to model autism spectrum disorder; they evidenced increased and decreased cooperative behaviors in Shank2 KO and Shank3 KO mice, respectively, which additionally correlated well with altered social dominance behaviors^14^.

In the above work, which is the only one using mice for mutualistic cooperation study, to our knowledge, the researchers assessed mice’s cooperative behavior with an automated cooperation test, an assay originally developed by Avi et al. in rats^15^. Comparing the performance between mice and rats in this assay, we found the efficacy (defined as the number of rewards divided by activity) in mice was only about half that in rats, which further reduced to¼ considering the mouse box is also half-long. Futhermore, the latency (defined as time to receive the first reward) in mice is one order of magnitude higher^14,15^. The low efficacy and high latency in mice make the assay insensitive for detecting subtle behavioral alterations. Therefore, although it is natural to consider borrowing behavior assays originally developed for rats, their performance in mice is often not optimal; de novo development of new assays for mice is necessary and beneficial.

### Experimental design

In this protocol, we established a new method to evaluate the cooperative behaviors of rodents including two models (‘altruistic cooperation’ and ‘mutualistic cooperation’). We use water as rewards to promote the cooperation of the animals. We set a location task for the subjects to get the rewards. In ‘altruistic cooperation’ models, one subject must sacrifice its own profit to execute the task for the reward given to the other. However, in ‘mutualistic cooperation’ model, two participants should finish the task at the same time to get two water rewards. The described protocol was assessed by different animal models considering the factors including gender, age, genetics and so on.

### Mouse models

In this experiment, we used different model mice to evaluate our new cooperative behavior test (CBT). Firstly, we used 2-month-old C57BL/6 male mice to establish our CBT successfully. However, we found that “mutualistic cooperation” model was suitable for mice whereas “altruistic cooperation” model was too difficult for them to finish the task. Thus, we use “mutualistic cooperation” model in our CBT sequentially. Secondly, in order to investigate whether gender or age has an effect on cooperation, we tested 2-month-old C57BL/6 male or female mice and C57BL/6 male mice of 2-to 9-month-old with CBT and found some difference between genders and ages (see anticipated results). Finally, we used two models with social interactive defection, APP/PS1 mice and chronic social defeat stress (CSDS) mice, to evaluate the reliability of our CBT.

### Setup of the apparatus

The CBT apparatus was designed as **Figure 1a** and **1b**. The CBT apparatus consist of two chambers. The size of upper chamber is 30 cm (length) × 20 cm (width) × 15 cm (height) and the lower is 30 cm (length) × 20 cm (width) × 2 cm (height). The two chambers are separated by a stainless-steel wire mesh. Rows of holes were punched in the walls for the airflow. For “mutualistic cooperation” model, two optoelectronic switches are set above the top wall 3 cm from the edge and the mirrors are set on the mesh of the corresponding location. For “altruistic cooperation” model, one optoelectronic switch is set above the top wall 3 cm from the edge and the mirrors are set on the mesh of the corresponding location. Water control valves are connected to the switches by wires and water bottles by pipes. For “mutualistic cooperation” model, two sippers are extended into the box just next to the mirrors from the water control valves on the top wall. While for the “altruistic cooperation” model, only one sipper is extended into the box 20 cm away for the mirror horizontally. The upper chamber is used for exploration of mice and the lower chamber is set as a drawer so that it is convenient to clean feces of mice.

**Figure 1.**
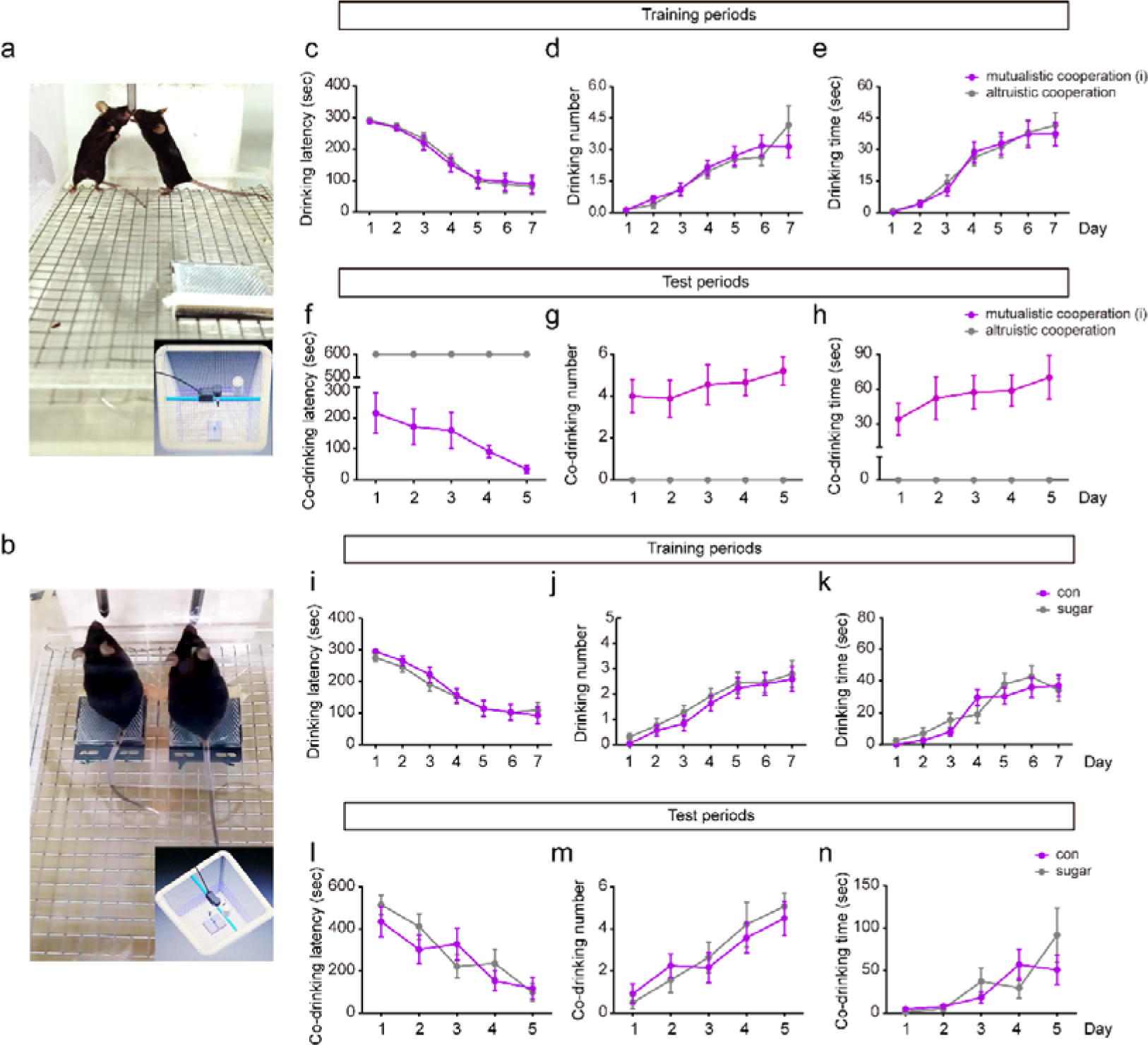
Establishment of cooperative behaviors test. (**a, b**) Diagram of our cooperative behaviors test apparatus, including “altruistic cooperation” model and “mutualistic cooperation” model. (**c-e**) The drinking latency, drinking number, and drinking time of mice in “mutualistic cooperation” and “altruistic cooperation” models during training periods. n = 23. For mutualistic cooperation: drinking latency, *F* _(6, 270)_ = 13.741, *p* < 0.001. drinking number, *F* _(6, 270)_ = 10.379, *p* < 0.001. drinking time, *F* _(6, 270)_ = 13.464, *p* < 0.001. For altruistic cooperation: drinking latency, *F* _(6, 270)_ = 18.792, *p* < 0.001. drinking number, *F* _(6, 270)_ = 10.671, *p* < 0.001. drinking time, *F* _(6, 270)_ = 14.321, *p* < 0.001, by ANOVA with Bonferroni’s post hoc test. (**f-h**) The co-drinking latency, co-drinking number, and co-drinking time of the pair of mice in “mutualistic cooperation” and “altruistic cooperation” models during test periods. n = 9. For mutualistic cooperation: co-drinking latency, *F* _(4, 180)_ = 2.173, *p* =0.09. co-drinking number, *F* _(4, 180)_ = 2.633, *p* > 0.05. co-drinking time, *F* _(4, 180)_ = 0.674, *p* > 0.05. For altruistic cooperation: co-drinking latency, *p* > 0.05. co-drinking time, *p* > 0.05. co-drinking time, *p* > 0.05, by ANOVA with Bonferroni’s post hoc test. (**i-k**) The drinking latency, drinking number, and drinking time of mice rewarded by regular or sugar implanted water in “mutualistic cooperation” and “altruistic cooperation” models during training periods. n = 25-30. (**l-n**) The co-drinking latency, co-drinking number, and co-drinking time of the pair of mice rewarded by regular or sugar implanted water in “mutualistic cooperation” and “altruistic cooperation” models during test periods. n = 12-14. The results are shown as the mean ± s.e.m. and tested by repeated-measures ANOVA with Bonferroni’s post hoc test.

### Water deprivation, habituation and tasks

Our CBT protocol is based on the water reward and therefore requires animals to undergo a water deprivation schedule to ensure that they are willing to do the task for obtaining water reward. As described in Step 3, water is removed from their home cages 8 h before training or testing. Habituation is also essential to reduce stress being introduced to the CBT apparatus. Also, this habituation can minimize the confounding effects of novelty on optoelectronic switches and sippers. In order to evaluate cooperative behaviors, we set up some tasks for the subjects. The animals must sit on the mirror to open the corresponding optoelectronic switch. For “altruistic cooperation” model, the mirror is away from sipper so that one animal should open the switch and the other get water reward (Here, we call it “task” in “altruistic cooperation” model). For “mutualistic cooperation” model, there are two switches and two sippers. The two animals should open the switches at the same time and therefore get two water rewards (Here, we call it “task” in “mutualistic cooperation” model). In this model, the distance of mirrors is long enough so that on animals cannot open the two switches simultaneously.

### CBT

Four parts, which comprises habituation, training, test and data analyses, are mainly involved in our CBT procedure. During habituation (Steps 1-2), animals are firstly moved to the test room for environment adaption. Then, they are transferred to the chambers for apparatus habituation 5 min a day for two days. At this stage, the animals have access to water ad libitum. During training stages (Steps 3-10), all animals are depriving of water 8 h before training. They are introduced to apparatus one by one to learn how to open the switch for water reward (Here, we call it “task” in training period). They are ready to take the test until all the animals have acquired the ability. This process will last for 5-7 days depending on the animals. During test stages (Step 11-18), a couple of animals are moved to the apparatus at the same time. During the next 10 min, they should learn to open the switch to get water for their companion in “altruistic cooperation” model or open the two switches simultaneously to get water for themselves. All the test process should be recorded by video system for analyses. All animals are given water at libitum in their home cages at the end of the test. This process lasts about 5 days. Finally, the latency to finish task, the number of the tasks accomplished and the time of their cooperation in every test are calculated to evaluate their cooperative ability (Step 19).

### Influences on heath or other behavior tests of subjects

We intend to assess whether long-term intermittent water deprivation affects the overall health and neuropsychiatric behaviors of mice. Therefore, before training or testing every day, we measure the weight of the animal. We also compared cognition associated or emotion associated abilities using Y maze, open field test, elevated plus maze and three chamber social test between animals with or without CBT. The results showed that there was no significant difference between two groups. However, the performance in Y maze and three chamber social test were slightly improved. This improvement may be due to that learning task every day enhanced the function of CNS.

### Alternative improvement of “mutualistic cooperation” model

The original vision of “mutualistic cooperation” model is that the switches are just beside the sippers. Thus, the animals could open the switches meanwhile get water rewards. However, this “cooperative behavior” could attribute partly to their desire for water separately. Therefore, we move the switches to the opposite of the sippers, and there are about 20 cm between switches and sippers so that the animals cannot complete tasks and get water reward meanwhile. This improved vision separates the willing to cooperate and the desire for water and could be accurate for cooperative behaviors studies. However, we use water restriction in our CBT and the desire for water is their motivation to cooperate. So that the two visions of “mutualistic cooperation” models can apply to monitor cooperative behaviors successfully. Notably, the task in the new vision is more difficult, so that it takes more time for the animals to acquire the ability to finish the tasks and the latency is longer in new vision than the older.

### Data collection and analyses

A video-tracking system is recommended because a video recording allows for the analyses by more than one experimenter in the following day. The camera can be set vertically or horizontally to the apparatus. Here, we recommend vertical way to monitor the animals. In our CBT, we mainly use three parameters to evaluate cooperative ability including latency, number of the tasks accomplished, time of drinking, and time of the cooperation. All the parameters are based on the finish of tasks. The tasks including opening the switch in training, opening the switch to get water for the other or opening switches simultaneously. The latency is defined as the time of finish the task firstly during training or test period. The number of the tasks accomplished is defined as the sum of numbers that they finish tasks during training or test. The time of drinking is defined as the sum of the time for drinking during training. The time of the cooperation is defined as the sum of time that the two tested mice are drinking water. Mindfully, different experiments should meet the same criterion of determining the finish of task.

## MATERIALS

### Reagents

Laboratory mice. Male or female wide-type or APP/PS1 mice in C57BL/6J or CD1 background, at different ages, can be used in this test. All experiments must receive approval from the relevant institutional review board and be conducted in accordance with local and national regulations.

Drinking water.

75% (vol/vol) ethanol for cleaning the equipment.

### Equipment

Cooperative test apparatus (made in-house; see Equipment setup), including cages, water pipes, optoelectronic switches (involving switches and mirrors), water control valves, wires and stainless-steel wire mesh that fit the cage dimensions.

Water bottles, equipped with standard stainless-steel sipper.

Electronic timers.

### Reagents setup

#### Mice

In our experiments, we used 8 to 10-week-old male and female C57BL/6J or CD1 mice, or 5-month-old WT and APP/PS1 mice in C57BL/6J background. Mice were housed in standard cages with a constant temperature (22°c ± 1°c) on a 12-h light-dark cycle (light 10:00-22:00). Food and water were given ad libitum except during the test. For cooperative test, mice were depriving of water for 8 hours twice a day.

! CAUTION All experiment carried out should be approved by the Institutional Animal Care and Use Committee of Nanjing Medical University.

### Equipment setup

#### CBT apparatus

The CBT apparatus is a rectangular box with a size of 30 cm (length) × 20 cm (width) × 17 cm (height). The apparatus was made of polyethylene material that is 0.4 cm thick and the walls were assembled by a super-bonding compound. The gate of the apparatus is on the side wall. Rows of holes were punched in the walls for the airflow. The optoelectronic switches are set above the top wall and the mirrors are set on the mesh of the corresponding location. Water control valves are connected to the switches by wires and water bottles by pipes. All these can be controlled by a main switch. Sippers are extended into the box for the drinking from the water control valves on the top wall. The apparatus is divided into two chambers by the stainless-steel wire mesh. The size of upper chamber is 30 cm (length) × 20 cm (width) × 15 cm (height), and the lower is 30 cm (length) × 20 cm (width) × 2 cm (height). Mice should be put into the upper chamber. The lower chamber is set as a drawer so that it is convenient to clean feces of mice.

### Experiment room

An air-conditioned room (22 ± 2 °C) is required for cooperative test. Also, the dim light is also needed. Experimenters should stay one meter away from the apparatus at least.

! CAUTION Avoid noise, light, and other environmental changes during the test.

### Data collection system

A video-tracking system is alternative (for instance, TSE Systems). These software packages need a computer system and a video camera located vertically to the apparatus. Otherwise, the data can be collected manually by experimenters who should be blinded to the groups. It is recommended to use a video-tracking system because a video recording allows for the analyses by more than one experimenter. ▲ CRITICAL The experimenter should have been trained to identify the cooperative behavior by a mouse so that the data can be valid and reproducible. If a single experimenter cannot finish the whole experiment, it is important to ensure that the same criteria must be met to identify a cooperative behavior.

## Procedure

### Preparation for training (day 1-day 3)

1. You should transport subjects to the behavior test room 3 days before test for acclimatization. You may begin handling and weighing the subjects during these days.

2. Apparatus adaptation. Clean up the apparatus with 75% (vol/vol) ethanol and wait them to be dry. Then place the subjects into the apparatus one by one with the main switch off once a day for 5 min. These should be done 2 days before the finish of acclimatization.

▲ CRITICAL The apparatus should be cleaned up between two consecutive adaptations.

### Training (day 4 – day 10)

3. Begin water restriction and maintain it throughout the whole test. It is recommended to restrict the water 8 h before the test. ! CAUTION Be sure to adhere to guidelines of governmental and institutional animal welfare.

4. Weigh each animal every day. ! CAUTION Avoid weight reduction of more than 5% per day or 85% of the whole test.

5. Set up the apparatus with the main switch on for the habituation of the subjects. Promise enough water in the bottles so that the subjects can get rewards if they finish tasks. For “mutualistic cooperation” models, one of the two optoelectronic switches should be made on during the training days so that one animal can finished the “cooperative” task and get water reward. ! CAUTION Choose one of the two optoelectronic switches on at liberty during every training to prevent the animals from habituating only one switch.

6. Move a single subject into the apparatus and allow 5 min to explore the apparatus and learn how to finish the task and gain rewards. It is recommended to use the video-tracking systems to detect the animals during this period if possible.

▲ CRITICAL If the subjects cannot fulfill a task during the training time, it is necessary to introduce them to make the switch on and get water rewards for at least 3 times.

▲ CRITICAL The times of training is important for the subjects to learn to use the equipment. The subjects will perform better and learn faster as the times of training increase every day. We recommend train the animals twice a day.

▲ CRITICAL We recommend to mark the tail of each animal with a permanent marker. So, it is convenient for you to distinguish the training status of individual animal.

7. After training termination, return animals to their respective home cages.

8. Clean up the switches, sippers, and chamber with 75% (vol/vol) ethanol and ensure them dry before the next training.

9. Repeat step 6-8 until all subjects have been trained.

10. Repeat step 3-9 twice a day for 7 days. ▲ CRITICAL Make sure all the subjects can learn to use the equipment and get water rewards until the finish of training days. Generally, five days are needed to get this point. You can extend the training periods appropriately.

### Test (day 11 – day 15)

11. Repeat step 3 for all test days.

12. Turn on the main switch and ensure that optoelectronic switches can work well. Start the video-tracking systems to monitor the animals successfully. Check if there is enough water in bottles and the water rewards can be given properly.

13. Clean up the switches, sippers, and chamber with 75% (vol/vol) ethanol and ensure them dry before the test.

14. Move a pair of subjects into the “altruistic cooperation”, “mutualistic cooperation” (i) or “mutualistic cooperation” (ii) apparatus and allow 10 min for them to get rewards cooperatively.

▲ CRITICAL Make sure that all the subjects have learned to use these models for rewards. If not, repeat the training sessions again.

▲ CRITICAL Three ways can be used to choose the pair of subjects. (i) Choose the two animals at random. (ii) Choose a matched pair of subjects every test. (iii) Use another animal (demonstrator) and match all the subjects to it successively. All these ways can be adapted for different purposes. Here, we recommend the second one for the normal test. ! CAUTION Do not use the same demonstrator in a consecutive test if you adopt the third way.

▲ CRITICAL Experimenters should stay at another room or at least 1 meter from the apparatus since the test is going on. So, we recommend a video-tracking system to record the performance of the subjects. It is useful for you to analyze the result at the same time of or after the test.

15. Return animals to their home cages after the tests is finished.

16. Clean up the switches, sippers, and chamber with 75% (vol/vol) ethanol and ensure them dry before the next test.

17. Make the main switch off and clean the bottles and sippers until all the subjects have been tested.

18. Repeat the test (step 11-17) once a day for 5 days.

## Data analyses (day 15)

19. Analyze the performance of subjects during the test as the following parameters: latency to get water rewards cooperatively, the number of the tasks accomplished and the time of their cooperation. ▲ CRITICAL Cooperation is defined that one subject stay on the mirror to open the optoelectronic switch and the other drink water at the same time in “altruistic cooperation” model or both subjects stay on the different mirrors to open the optoelectronic switches and get water rewards meanwhile in “mutualistic cooperation” model.

### Troubleshooting

| Step | Problem | Possible reasons | Solution |
| --- | --- | --- | --- |
| 3 and 4 | Significant weight loss | Water restriction | Give water appropriately |
| 5 and 12 | Leaking or blocking of tubes | Broken tubes | Replace with new tubes and keep air tight |
| 10 | A mouse does not reach the criterion | Stress | Exclude the mouse from the analysis |
|  | Most of the mice do not reach the criterion | (i) Ineffective water restriction | Restrict water for a long time |
|  |  | (ii) Stress | Check the experimental conditions: temperature, humidity, luminosity, noise, or odors related to the experimenter |
| 14 | Most of the mice do not finish the tasks cooperatively | (i) Ineffective water restriction | Restrict water for a long time |
|  |  | (ii) Stress | Check the experimental conditions: temperature, humidity, luminosity, noise, or odors related to the experimenter |

### Timing

Step 1, acclimatization for behavior test environment and handling: 1 day, 3 min per animal

Step 2, apparatus adaptation: 2 days, 5 min per animal

Step 3-10, training: 7 days, 5 min per animal

Step 11-18, test: 5 days, 10 min per animal

Step 19, data analyses: 1 day

### Anticipated results

Three parameters are used here to evaluate the cooperative ability in our CBT: the latency for the first successful co-drinking, the times of co-drinking in each trial, and the cumulated time of co-drinking in each trial. The latency is more reflective of their cooperation ability, because their desire for water should be highest at the beginning; the co-drinking number and the cumulated time are more reflective of the amount of water they required during each trial.

To demonstrate the capability of our CBT in measuring mice’s cooperative ability, we introduced 2-month-old male mice into our apparatus. After the 7 days of individual training, all mice learned to accomplish the task for water rewards, as reflected by the declining latency curve (**Fig. 1c-e & Supplement Video 1**). Next, we moved those mice to the test setup. In the “mutualistic cooperation (i)” model, through the testing phase, we observed constantly declined latency with significant improvement among trials, while the co-drinking number and the cumulated time showed rising trends, but without statistical difference (**Fig. 1f-h & Supplement Video 2**). In the “altruistic cooperation” model, however, all three parameters remained unchanged throughout (**Fig. 1f-h & Supplement Video 3**). Therefore, these results demonstrate that mice in this setup demonstrate mutualistic but not altruistic cooperative behavior. Next, we kept only the “mutualistic cooperation” model to evaluate mice’s mutualistic cooperative performance. We also tried the 1% sucrose solution instead of the regular water as a reward. We expected to a shorted training and testing phase. However, although the latencies further decreased slightly, we didn’t see a statistical difference (**Fig. 1i-n**). Therefore, the following experiments were still carried out with regular drinking water.

Next, the weight of mice and general behavior tests were detected to evaluate the influence of our CBT on the subjects. The weight of mice with or without the CBT showed no difference (**Fi. 2a**). The anxiety or depression related behaviors were also unchanged detected by open field test or elevated plus maze between two groups (**Fig. 2b-e**). There was also no difference in memory associated test such as Y maze (**Fig. 2f-2g**). In the novel object recognition test, ratio of time spent in sniffing novel object in total sniff was slightly elevated but no difference (**Fig. 2h**). These could be attribute to that training and test every day in our CBT might have an improving effect on thememory and cognition, which was in accordance with that social interactive task may delay dementia^13^.

**Figure 2.**
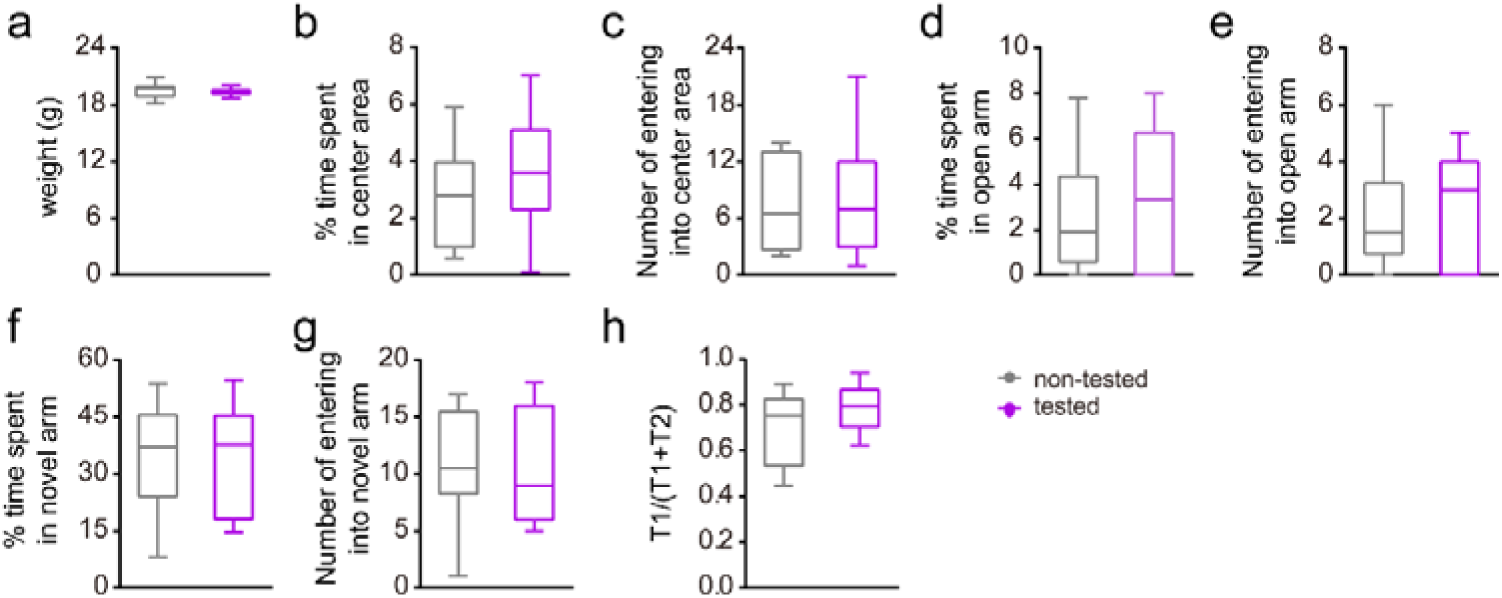
Our cooperative behavior test did not influence the weight and other behaviors performance of the mice. (**a**) The weight of mice with or without our cooperative behavior test. (**b, c**) The percentage of time spent in center area and the number of entering into the center area in open field of mice with or without our cooperative behavior test. (**d, e**) The percentage of time spent in open arm and the number of entering into the open arm in elevated plus maze of mice with or without our cooperative behavior test. (**f, g**) The percentage of time spent in novel arm and the number of entering into the novel arm in Y maze of mice with or without our cooperative behavior test. (**h**) The ratio of time spent in sniffing novel object in total sniff during novel object recognition test of mice with or without our cooperative behavior test. n = 10-19. The results are shown as the mean ± s.e.m. and tested by Student’s t-test.

In order to reveal whether gender or age have an influence on cooperative ability or not, 2-month-old male or female mice or 2 to 9-month-old mice were test in our CBT. The results showed that female mice have cooperative behavior in our CBT as well (**Fig. 3a-f**). However, female mice spent more time to learn to finish the task during training period and learn cooperation during test period (**Fig. 3d**). Moreover, the number and time of cooperation in female mice were decreased compared with male mice (**Fig. 3e-f**). This difference may be due to their sensitivity to scare. In aspect of age, there were no difference in latency, number, or time of cooperation between 2, 5 or 9-month-old male mice (**Fig. 3g-l**). But the social capacity of old mice remains to be evaluated.

**Figure 3.**
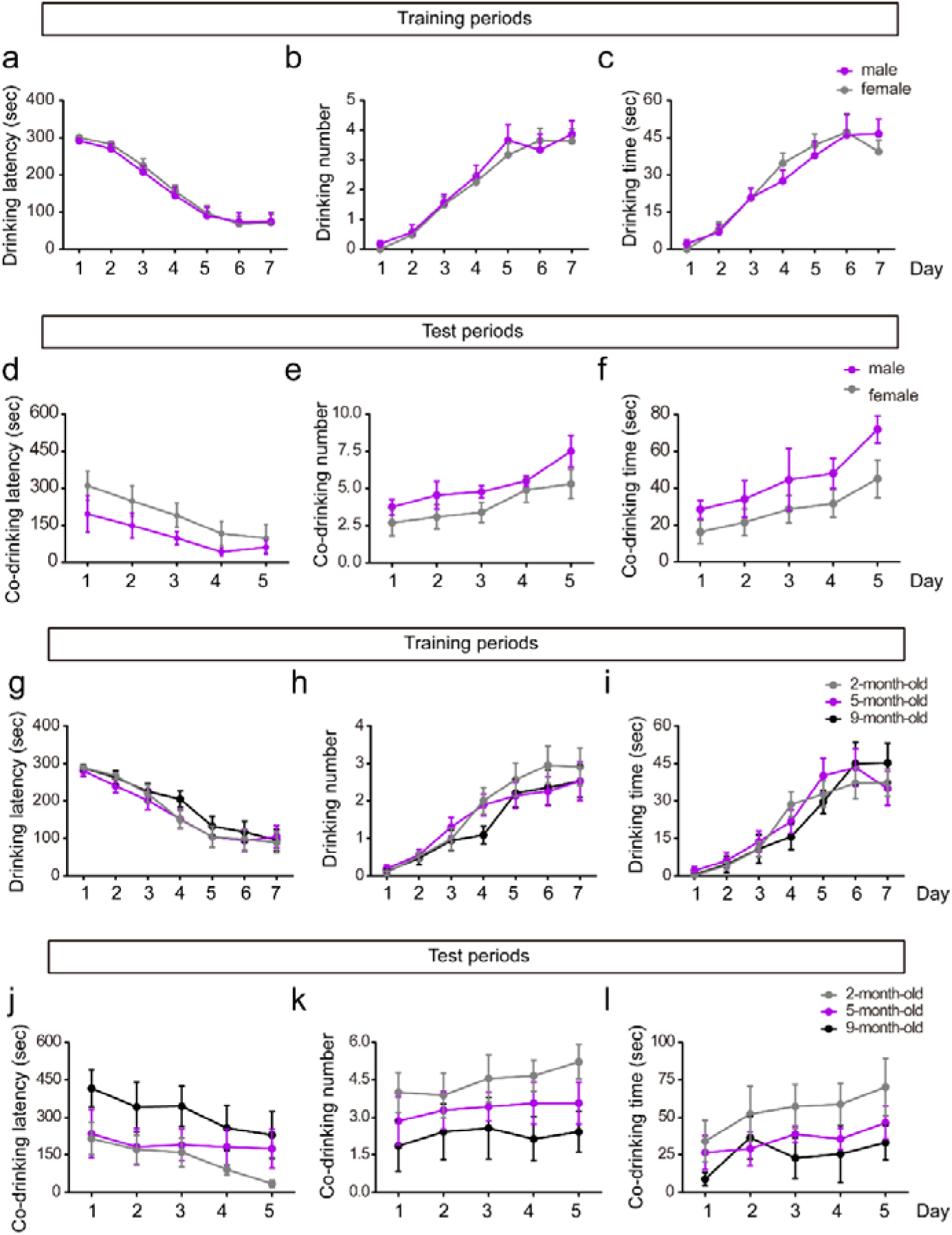
Detection of cooperative behaviors in mice with different genders and ages by our models. (**a-c**) The drinking latency, drinking number, and drinking time of male and female mice in “mutualistic cooperation” and “altruistic cooperation” models during training periods. n = 22-23. (**d-f**) The co-drinking latency, co-drinking number, and co-drinking time of male and female mice in “mutualistic cooperation” and “altruistic cooperation” models during test periods. n = 8-10. (**g-i**) The drinking latency, drinking number, and drinking time of 2, 5, 9-month-old mice in “mutualistic cooperation” and “altruistic cooperation” models during training periods. n = 17-23. (**d-f**) The co-drinking latency, co-drinking number, and co-drinking time of 2, 5, 9-month-old mice in “mutualistic cooperation” and “altruistic cooperation” models during test periods. n = 7-9. The results are shown as the mean ± s.e.m. and tested by repeated-measures ANOVA with Bonferroni’s post hoc test.

Then, two other mice models, APP/PS1 mice and CSDS mice were used to qualify the feasibility of our CBT. Previously studies have reported that the social interactive ability was impaired in 5-month-old APP/PS1 mice or CSDS mice and social interactive ability was fundamental for cooperation behaviors [14-16]. This result revealed that our CBT was applied for testing cooperative behaviors in a wide range of mice models. Taken together, the set of procedures and results above provide a reliable and useful method to evaluate cooperative behaviors (**Fig. 4a-l**).

**Figure 4.**
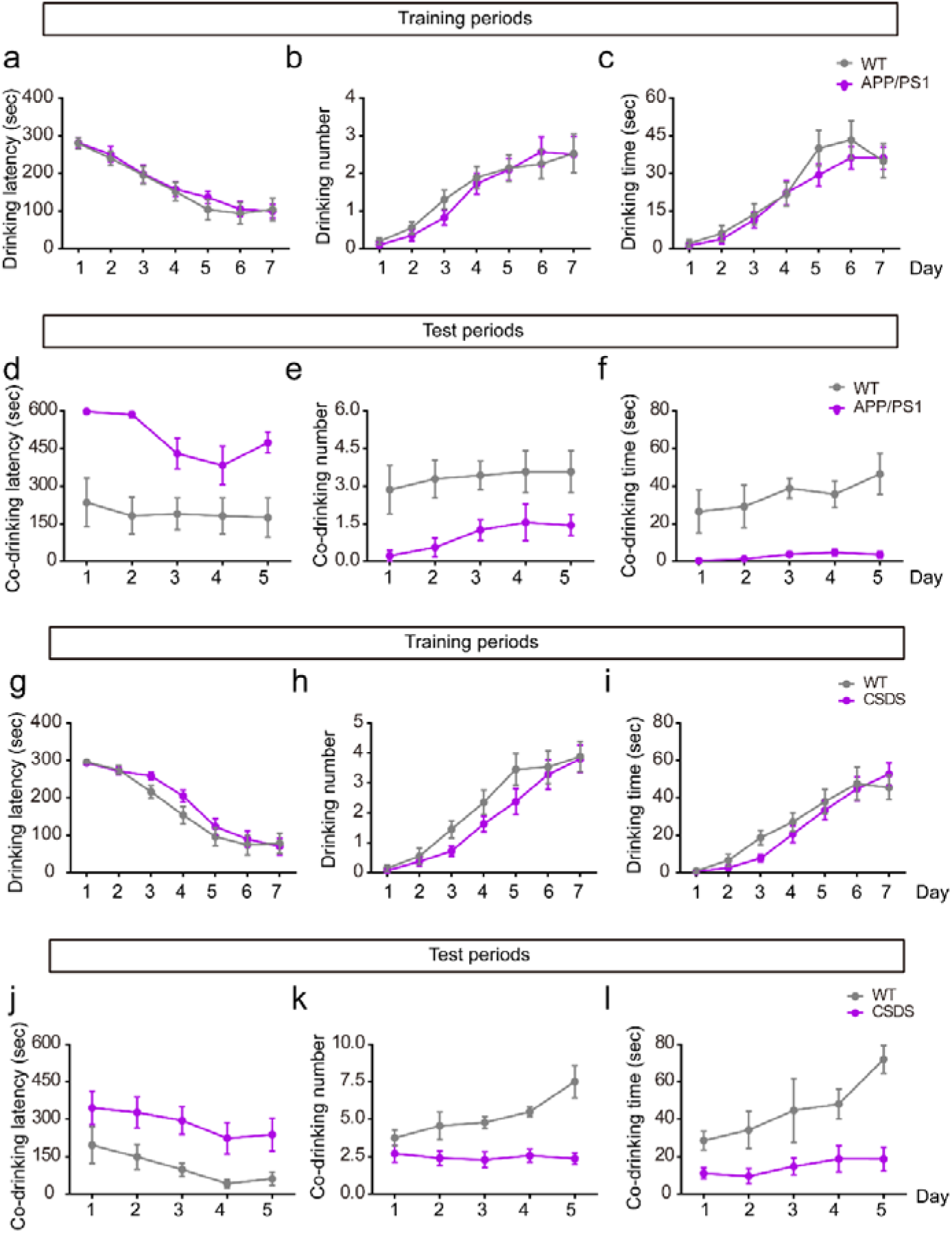
Detection of cooperative behaviors in social ability impaired APP/PS1 and CSDS mice by our models. (**a-c**) The drinking latency, drinking number, and drinking time of WT and APP/PS1 mice in “mutualistic cooperation” and “altruistic cooperation” models during training periods. n = 18-20. (**d-f**) The co-drinking latency, co-drinking number, and co-drinking time of WT and APP/PS1 mice in “mutualistic cooperation” and “altruistic cooperation” models during test periods. n=7-9. For drinking latency, *F* _(1, 375)_ = 28.028, *p* < 0.001. For drinking number, *F* _(1, 375)_ = 64.989, *p* < 0.001. For drinking time, *F* _(1, 375)_ = 69.421, *p* < 0.001. (**g-i**) The drinking latency, drinking number, and drinking time of WT and CSDS mice in “mutualistic cooperation” and “altruistic cooperation” models during training periods. n = 40. (**d-f**) The co-drinking latency, co-drinking number, and co-drinking time of WT and CSDS mice in “mutualistic cooperation” and “altruistic cooperation” models during test periods. n = 7-10. For co-drinking latency, *F* _(1, 460)_ = 20.532, *p* < 0.01. For co-drinking number, *F* _(1, 460)_ = 29.301, *p* < 0.001. For co-drinking time, *F* _(1, 460)_ = 14.534, *p* < 0.01. The results are shown as the mean ± s.e.m. and tested by repeated-measures ANOVA with Bonferroni’s post hoc test.

Finally, to exclude the possibility that the pair of mice completed the tasks due to their desire for water separately, but not cooperatively, the original “mutualistic cooperation (i)” and the improved “mutualistic cooperation (ii)” models were used. The mice could finish the cooperative task as well (**Fig. 5a-e**), which proved that the design of our CBT was useful to test cooperative behaviors, independently on where the switches was. As expected, the tasks in “mutualistic cooperation (ii)” were more difficult than that in “mutualistic cooperation (i)”, so the latency were extended and the cooperative behaviors were decreased in “mutualistic cooperation (ii)” model (**Fig. 5d**). Then, we compared the difference between the previous cooperative test model and ours. In the previous model, the participants are separated by a grille partition. Although they can see, hear, and smell each other, most of physical contact are deprived. We used the similar grille partition in our CBT to separate the mice and detect cooperative behavior subsequently. The mice separated by partition spent more time to learn to finish the task during the training period. In the test period, the latency was extended and the number and time of cooperation were also decreased significantly (**Fig. 5f-k**). The cooperative behavior needs more than sight and smell, so our CBT provided a better place for the mice to touch to finish the cooperative tasks. However, the mechanism of this cooperative behavior requires further investigation.

**Figure 5.**
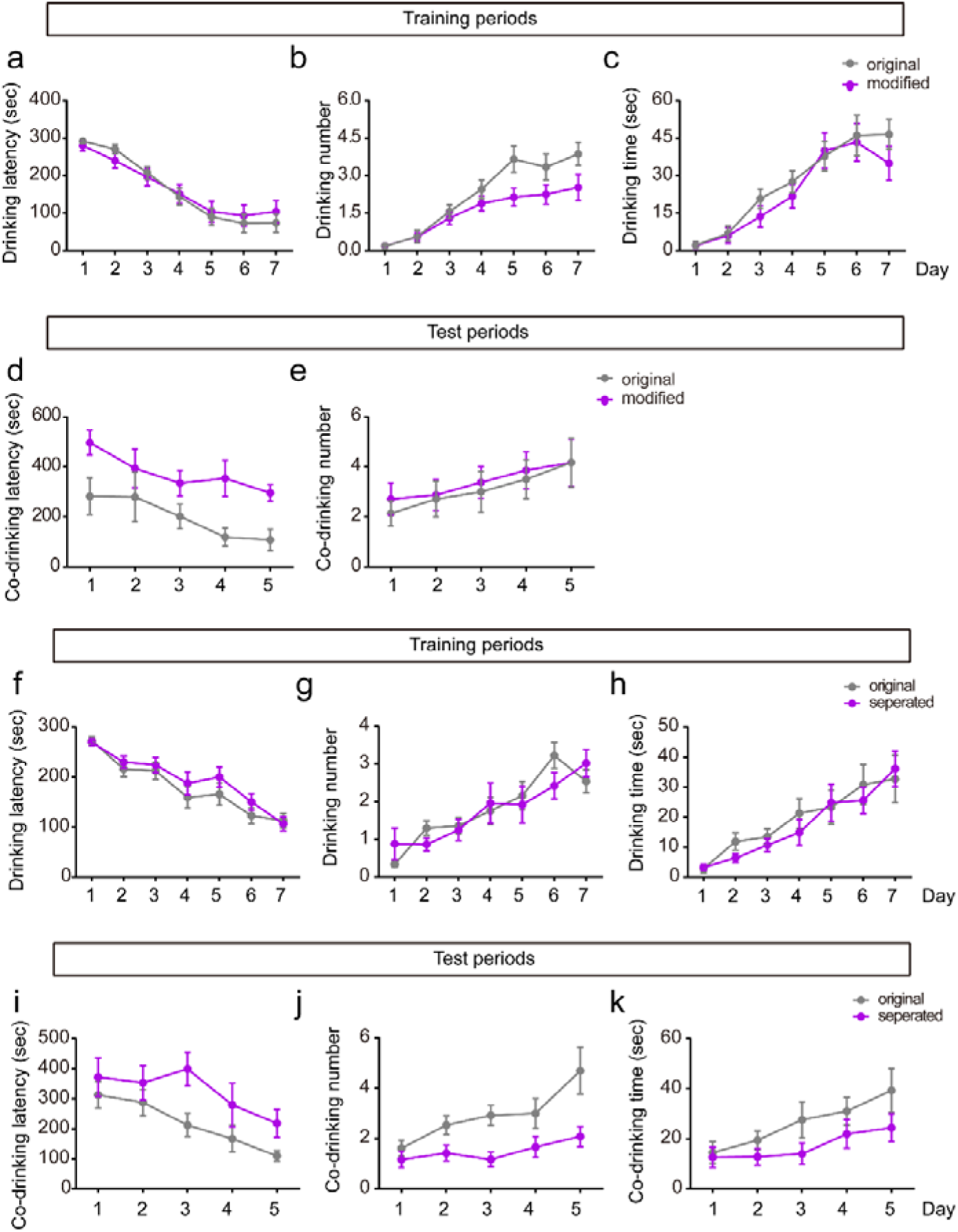
The modified vision of our cooperative behavior test. (**a-c**) The place of the switches did not influence the latency to cooperation and the number and time of cooperation in “mutualistic cooperation” models during training periods. n = 19-22. (**d-e**) The latency to cooperation was increased in modified “mutualistic cooperation (ii)” models during test periods. n = 6-8. For co-drinking latency, *F* _(1, 375)_ = 12.712, *p* < 0.05. For co-drinking number of cooperation, *F* _(1, 375)_ = 5.4, *p* > 0.05. For co-drinking time, *F* _(1, 375)_ = 69.421, *p* < 0.001. (**f-h**) The drinking latency, drinking number, and drinking time of the mice in models where they can touch each other or not during training periods. n = 20-30. (**i-k**) The co-drinking latency, co-drinking number, and co-drinking time of the mice in models where they can touch each other or not during test periods. n = 12-13. For co-drinking latency, *F* _(1, 625)_ = 4.198, *p* =0.052. For co-drinking number, *F* _(1, 625)_ = 10.370, *p* < 0.01. For co-drinking time, *F* _(1, 625)_ = 3.351, *p* = 0.08. The results are shown as the mean ± s.e.m. and tested by repeated-measures ANOVA with Bonferroni’s post hoc test.

## ACKNOWLEDGEMENTS

This work was supported by the grants from by the National Natural Science Foundation of China (81801378 and 81871117).

## AUTHOR CONTRIBUTIONS

M.X. and C.S. designed the original experiment. W.F. and Y.Z. designed the testing apparatus. W.F., Y.Z., Z.W., T.W., Y.P. and Y.Z. performed behavioral tests and the analyses. W.F. and Y.Z. did image typesetting. W.F., Y.Z. and H.H. wrote the primary version of the manuscript. M.X. and C.S. revised the manuscript.

